# A free classified advertising website is a high-risk unregulated source of domestic wild boar, pot-bellied pigs, and their hybrids with domestic swine in Canada: Implications for African Swine Fever Risk Preparedness

**DOI:** 10.1101/2022.03.14.483774

**Authors:** Amanda M. MacDonald, Ryan K. Brook

## Abstract

Invasive wild pigs are a global problem and are established in large areas across Canada. Domestic European wild boar were first introduced to Canadian farms in the 1980s, and their escape and intentional release have since led to free-ranging populations, which have expanded rapidly in numbers and range. Wild pigs are associated with widespread damage to crops and natural environments, as well as risks of disease transmission. African Swine Fever (ASF) is a highly contagious viral disease of global concern, that can spread between domestic and free-ranging swine and cause catastrophic economic impacts. As ASF preparedness requires a detailed understanding of the distribution and movements of all pig types, this paper aimed to characterize an unregulated ‘grey’ swine market in Canada to understand risks to domestic swine production and potential contributions to free-ranging wild pig populations. Kijiji.ca is a free Canadian internet classified advertising service, almost exclusively cash-based with no receipts and few, if any, records kept of transactions. We monitored Kijiji for sales of domestic wild boar, pot-bellied pigs, and their hybrids across Canada over two months from April 28 - June 30, 2021. Data was collected from all advertisements, including how the seller labelled pigs’ breed, age, sex, number for sale, sexual intactness, presence of tattoos and/or ear tags, as well as the date and location of the posting. Locations were mapped and compared to the spatial distribution of existing free-ranging wild pigs in Canada to identify areas they were likely to facilitate new populations or potentially supplement and genetically diversify existing populations. We identified 151 advertisements on Kijiji, the most coming from Ontario (34.4%; n = 52) and Alberta (29.1%; n = 44), followed by Saskatchewan (11.3%; n = 17), British Columbia (10.6%; n = 16), Nova Scotia (7.3%; n = 11), New Brunswick (4.0%; n = 6), Manitoba (2.0%; n = 3), Quebec (0.7%; n = 1), and Prince Edward Island (0.7%; n = 1). No listings were observed in Newfoundland and Labrador, or Yukon, Nunavut and Northwest Territories. Overall, 62 (41%) of advertisements were within watersheds with existing wild pigs that may contribute to supplementing additional swine, and 89 (59%) were within watersheds where wild pigs have not yet been identified. African swine fever preparedness should include policies and action to address this unmonitored and unregulated sale of swine in Canada, including ear tags, tattoos, genetic analysis, and mandatory reporting.

## 1. Introduction

There are no native pigs (*Sus scrofa*) in Canada (Banfield, 1974; Aschim and Brook, 2019). Since being introduced in 1598, Canada has become one of the largest domestic pork producers in the world, with approximately 14 million animals spread out over 7,600 farms (Blair, 2016; Statistics Canada, 2021a, 2021b). Domestic European wild boar were introduced to most provinces in Canada in the 1980s and 1990s to increase livestock diversity on farms, and more recently the Yukon Territory (Brook and van Beest, 2014; Michel et al., 2017; Aschim and Brook, 2019). These domestic animals were cross bred with various domestic pig breeds to produce wild boar-domestic pig hybrids (Michel et al., 2017). Vietnamese pot-bellied pigs were also introduced to Canada in the 1980s (Porter et al., 2016). These pigs were normally raised on small farms and are primarily sold through the pet trade. Through escapes and intentional releases of domestic wild boar, domestic swine, pot-bellied pigs, and hybrids of these, free-ranging populations of invasive wild pigs (*Sus scrofa*) have been the sole source of free-ranging wild pigs and they have become firmly established across large areas of the western Prairie Provinces of Alberta, Saskatchewan, and Manitoba (Aschim and Brook, 2019). All members of the same species and referred to as feral swine, wild hogs, feral hogs, feral pigs (Keiter et al. 2016), razorbacks, and incorrectly as ‘wild boar’, free-ranging invasive wild pigs in Canada have been detected in all provinces except Atlantic Canada, though there is not clear evidence of established populations outside of the Prairie Provinces in western Canada (Aschim and Brook, 2019).

Invasive wild pigs are continuing to expand exponentially in number and distribution in Canada and the occurrences occupy an area larger than 1 million km^2^ (Aschim and Brook, 2019, Brook unpublished data, 2022). Wild pigs are one of the most invasive and destructive species globally, causing widespread damage to agricultural crops and the environment, at an often cited and likely conservative cost estimate of US$1.5B/yr in the USA, which was increased to US$2.5B/yr in 2021 (Pimental, 2007; Strickland et al., 2020; USDA, 2021). Wild and domestic pigs also may act as reservoirs for diseases and pathogens that can threaten other livestock, wildlife, pets, and humans (Wyckoff et al., 2009). African swine fever (ASF; Whiting, 2003), pseudorabies (Boadella et al., 2012; Gaskamp et al., 2016), swine brucellosis (Pedersen et al., 2012), porcine reproductive and respiratory syndrome (Albina et al., 2000), and porcine epidemic diarrhea (Brazeau, 2019) are diseases that could have significant economic impacts on Canada’s pork industry if they became prevalent in commercial swine. *Trichinella, Escherichia coli, Salmonella* and *Leptospira* (Hampton et al., 2006; Barrios-Garcia and Ballari, 2012; Miller et al., 2017) are pathogens of wild pigs that to cause disease in wildlife and humans. Lastly, with their invasive success and presence at the human-wildlife-livestock interfaces, wild pigs also have the potential to be significant reservoirs and vectors of antimicrobial resistant bacteria (Torres et al., 2018).

African swine fever is an extremely infectious, viral haemorrhagic disease of domestic and free-ranging pigs for which there is no commercial vaccine or therapeutic treatment currently available (Guinat et al., 2014; Pollock et al., 2021). Transmission of ASF is often through direct contact between uninfected and infected domestic and/or wild pigs with high morbidity and up to 100% mortality, (Sánchez-Vizcaíno et al., 2019; Corn and Yabsley, 2020). African swine fever has never been detected in Canada or the United States, however; early detection and preparedness is critical in the event of an outbreak and requires a detailed understanding of the distribution and movements of all types of swine throughout North America (Sánchez-Vizcaíno et al. 2019). Although invasive wild pig numbers are rising out of control in Canada, no testing or surveillance programs are in place to detect occurrences of African swine fever in their populations. Moreover, research on small scale backyard domestic pig or ‘exotic’ pig farms in Canada is absent.

Animals are bought and sold for pets and meat production through numerous channels, for example at shelters or rescues, auctions, and livestock markets. More recently, the internet has become a major avenue for sales of various breeds of swine, for example, domestic wild boar, domestic pigs, and hybrids of both, which in turn creates a gateway to drive the introduction and spread of non-native and invasive pigs and establishing numerous small scale swine farms (Duggan et al., 2006; Keller and Lodge, 2009; Chucholl, 2013). While most contemporary research has focused on the ecology and management of invasive wild pigs, little attention has been given to the number and types of pigs for sale on the internet and the risks they pose to augmenting existing feral pig colonies or creating new ones and their potential for spreading diseases such as African swine fever within and between domestic swine systems.

The objectives of this research were to: 1) characterize the spatial distribution, number and type of domestic wild boar, pot-bellied pigs, and their hybrids for sale in Canada on www.kijiji.ca, 2) identify spatial overlap in online sales and existing wild pig distribution in Canada, and 3) consider the implications of our findings in the context of ASF risk. This information will inform managers and regulators of the nature of sale of swine on the internet and will highlight previously unexamined anthropogenic sources of invasive wild pigs into the environment in Canada. These insights are critical to Canada’s efforts to optimize ASF preparedness and response in the event ASF did arrive in Canada.

## 2. Materials and methods

We chose to examine the online classified advertising website Kijiji (www.kijiji.ca) as it was the most popular online classified service in Canada at the time of this study and because we knew that private sales of all livestock was banned by Facebook in 2019. We monitored the Kijiji page over two months from April 28 2021, through to June 30 2021, for domestic wild boar, pot-bellied pigs, and their hybrids. The terms “boar”, “wild boar”, “wild pig”, “potbelly”, “potbellie”, “pot-belly”, and “pot-bellie” were searched under “all categories” with location set to “Canada”. Screenshots of each advertisement (and all photographs when provided) were saved for subsequent data collection. For each advertisement only publicly displayed information was recorded, such as the date of posting and the location of the seller based on reported postal code and nearest city or town. When provided (in seller description), data on breed, age (Merta et al., 2015), sex, presence of tattoos and/or ear tags, number of animals, and information on sexual intactness was also collected. We tested for differences in the count of advertisements in each province (standardized by size) compared to a random distribution using SPSS 28.

Previous distribution maps of free-ranging wild pigs in Canada (Aschim and Brook, 2019) were updated following the same methodology to include all reports and occurrences of wild pigs collected up to and including February 2022. ArcGIS was used to map out the locations of advertised Kijiji pig sales and compared to the current wild-pig distribution to identify if there were regions where pig sales overlapped with the existing wild pig range. Mapping of wild pigs was approved by the Behavioral Research Ethics Board at the University of Saskatchewan (BEH no. 15-155). Collection of data from Kijiji was exempt from ethical review by the University of Saskatchewan as it used only publicly available data and specific details from the advertisements were kept confidential and locations were mapped at coarse scales.

## 3. Results and Discussion

A total of 151 classified ads were identified between April 28 and June 30, 2021, as selling domestic swine, including wild boar, wild boar x domestic pig hybrids, pot-bellied pigs, and pot-bellied x wild boar hybrids (Fig.1). These pigs were further categorized into six breeds, the potbelly pig (i.e., mini, mini-Vietnamese, and Dwarf), mini pig (i.e., Juliana, American mini, teacup), wild boar, Kunekune, Ossabaw, and heritage pigs (i.e., Hampshire, Mangalitsa, Tamworth, Berkshire, Yorkshire, Landrace, Spotted, Hereford, Large Black, and Duroc). There were nine Canadian provinces with ads selling pigs, British Columbia, Alberta, Saskatchewan, Manitoba, Ontario, Quebec, New Brunswick, Nova Scotia, and Prince Edward Island (Fig. 2). No advertisements were observed from Newfoundland and Labrador or any of Canada’s three Territories. Data for the number of advertisements per province and breed are summarized in Table 1 and mapped by reported breed (Fig. 2). Distribution of advertisements within provinces standardized by province size were significantly different from random (*X*^2^ 6 = 2.1, *p* = <0.001), with occurrences ranked from highest to lowest occurring in AB (34%), ON (24%), Atlantic Canada (17%), SK (14%), BC (8%), MB (2%), and QC (<1%).

**Table 1.**
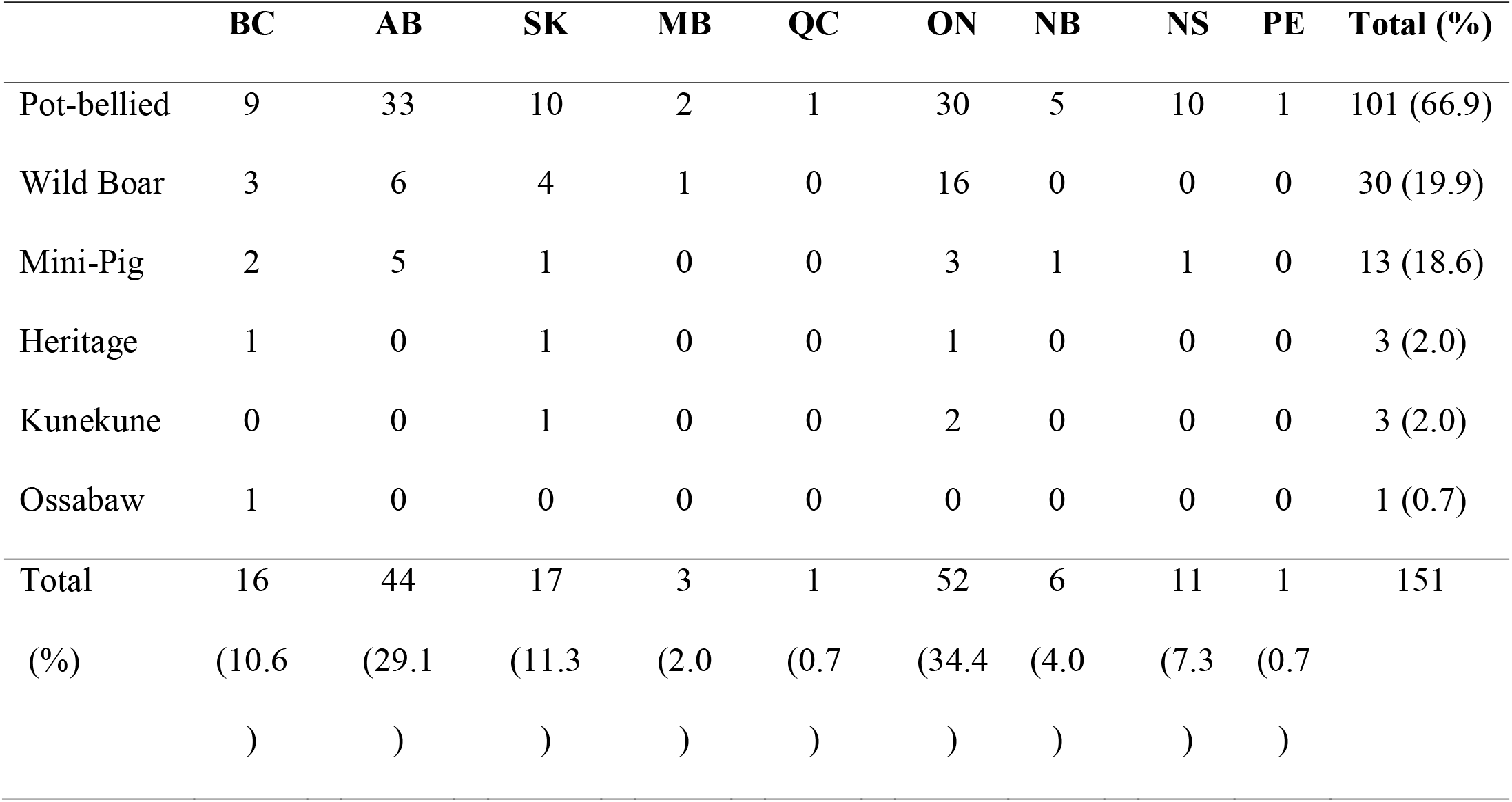
Summary of the provinces where advertisements for sale of domestic wild boar, pot-bellied pigs and hybrids with domestic pigs that were posted on the online sales website www.kijiji.ca between April 28 2021 to June 30 2021.

**Fig. 1.**
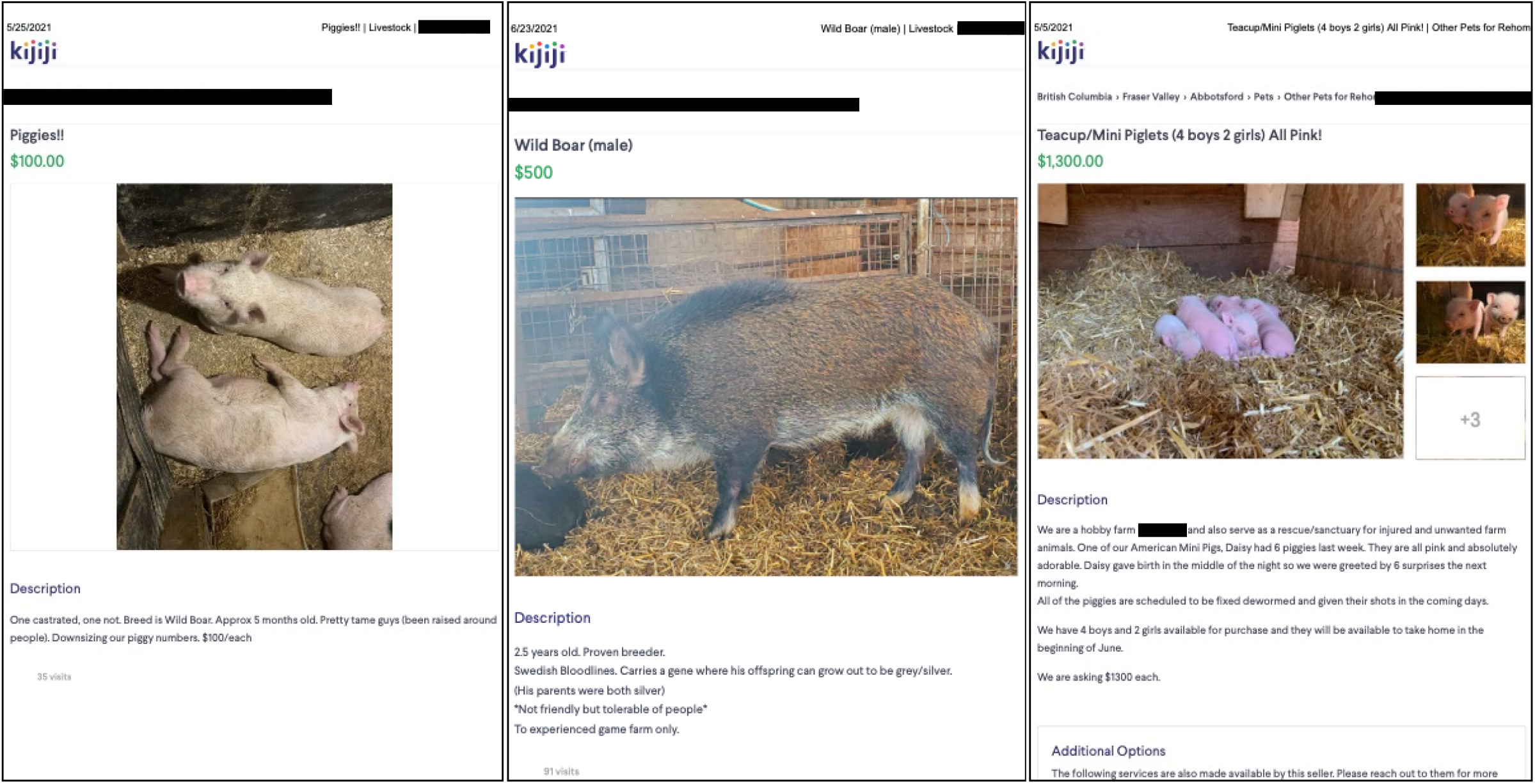
Screenshots of advertisements for sale of domestic wild boar, pot-bellied pigs and hybrids with domestic pigs that were posted on the online sales website www.kijiji.ca between April 28, 2021, to June 30, 2021. Specific details are blocked out to maintain confidentiality of individual operations.

**Fig. 2.**
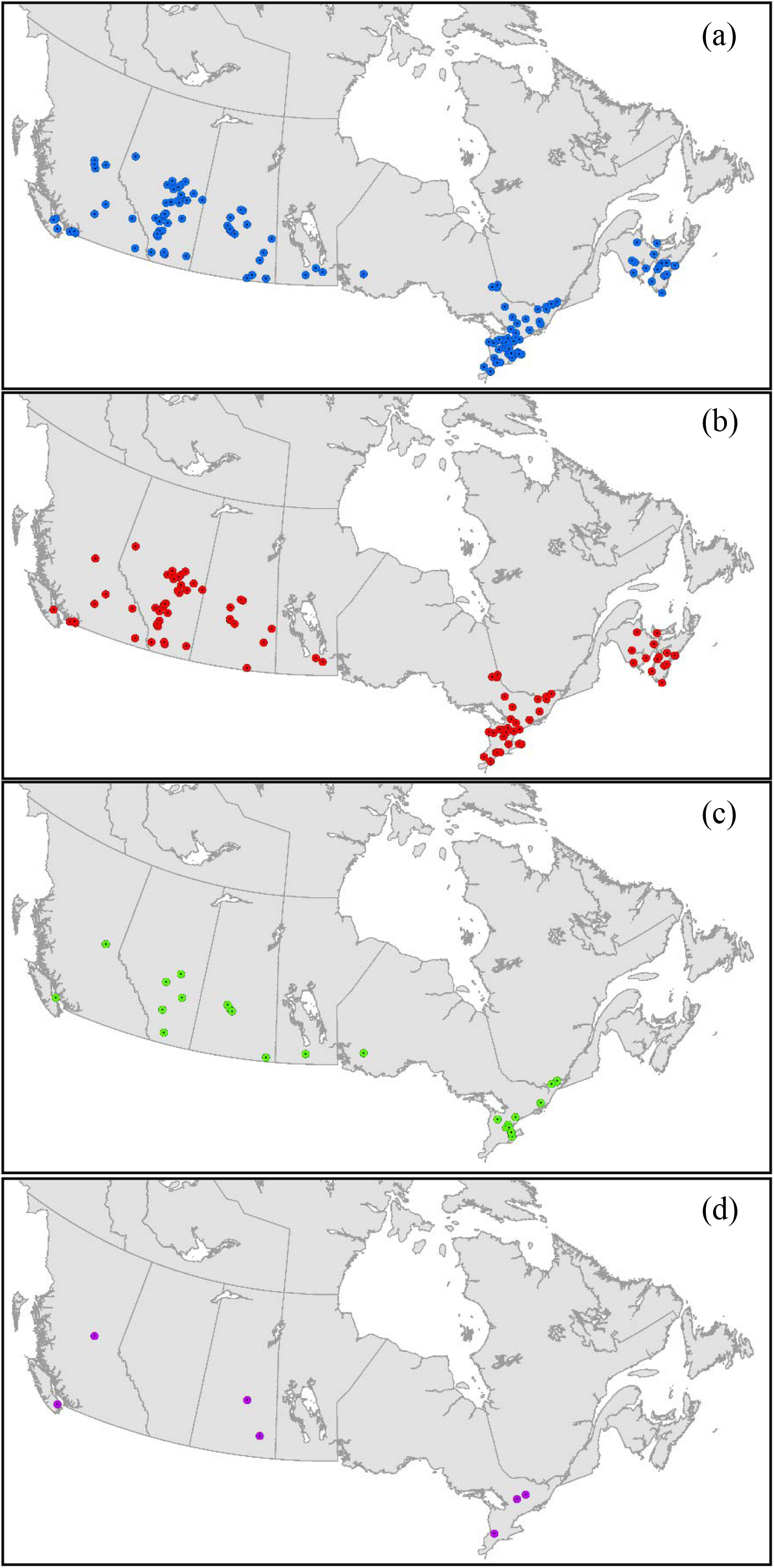
Map of Canada with (a) locations of all pig sales observed on Kijiji between April 28 - June 30, 2021, (b) locations of pot belly and mini-pig listings, (c) wild boar listings, and (d) locations of all other breeds i.e., heritage, Kunukune, and Ossabaw pigs.

In terms of age, most listings had piglets for sale (58.9%; n = 89), followed by adults (10.6%; n = 16), and yearlings (6.6%, n = 10). Eight (5.3%) ads offered pigs of various ages, and in 28 (18.5%) age was not specified. Offers of pigs in groups including both sexes were listed most often (34.4%; n = 52), males in 36 (23.8%) ads, and females in 32 (21.2%). Sex was not provided in 31 (20.5%) advertisements. Only four (2.6%) sellers reported pigs being tattooed or tagged, and no information was given by 147 (97.4%). The number of pigs for sale per ad ranged from 1-50 individuals. Fifty-seven (37.7%) ads reported sexually intact pigs for sale, 11 (7.3%) listed pigs that were not intact, and 83 (55%) did not specify.

Within Canada we documented 55,454 confirmed occurrences of free-ranging invasive wild pigs occupying 1,931 level 9 watersheds (Fig. 3). Overall, 62 (41%) of the Kijiji advertisements we documented were within watersheds with existing occurrences of wild pigs and may contribute to supplementing additional swine, while 89 (59%) were within watersheds where wild pigs have not been identified yet (Fig. 3). Of those locations not within wild pig established range, the wild boar types for sale were in British Columbia (15%), Alberta (31%), and Ontario (54%). However further research is needed to determine the distances that these swine are transported from the farms where they are purchased and what their fates are, including how many escaped or were released and/or escaped to the wild. Domestic and free-ranging wild pigs have a broad global distribution across all continents except Antarctica and continue to expand into new areas at rapid rates (Robinson et al., 2014; Lewis et al., 2017). In Canada, numerous types of domestic swine are held for meat production, high-fence shoot operations where people pay to harvest a wild pig inside a fenced compound, and as pets. Escaped or intentionally released domestic swine, including wild boar and their hybrids, have been the primary and likely only source of free-ranging wild pigs in Canada (Brook and van Beest, 2014; Koen et al., 2021). These invasive pigs are associated with considerable negative ecological and socio-economic impacts. Despite this, the number and types of domestic pigs available for sale on the internet, and their potential as a source of invasive wild pigs in Canada, has not been addressed.

**Fig. 3.**
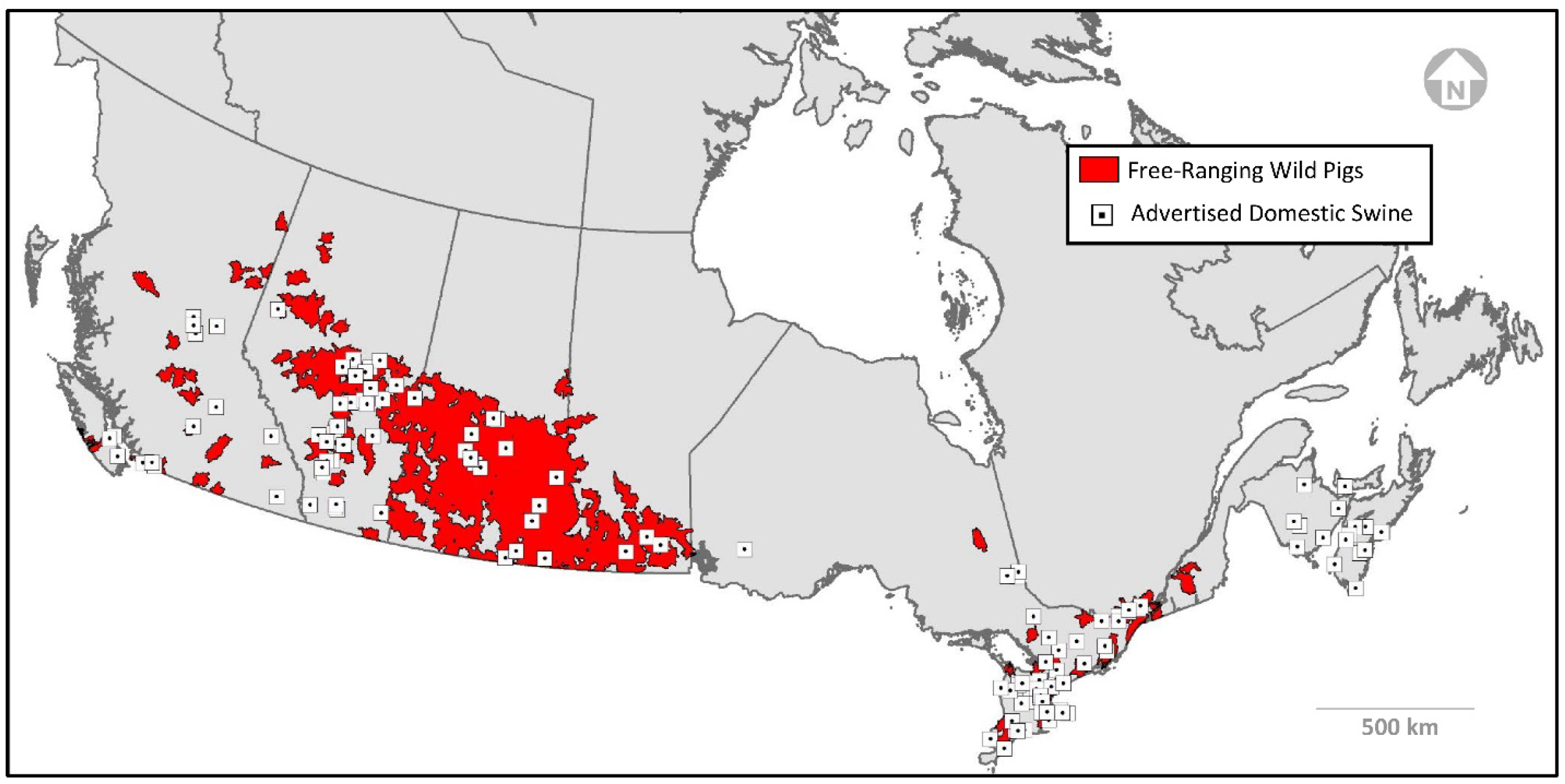
Spatial distribution of level nine watersheds that have free-ranging invasive wild pig occurrences (1995-2022) and locations of sales of domestic wild boar, pot-bellied pigs and hybrids with domestic pigs that were posted on the online sales website www.kijiji.ca between April 28 2021 to June 30 2021.

The website Kijiji.ca was the main source of information used in this study. Established in 2005, Kijiji is currently the most popular online classified service in Canada, functioning as a free network that is searchable by keywords (Kijiji, 2017). It is important to note here that sales and purchases are typically cash based with no receipts and no records kept of the transactions. Indeed, a core function of the website is to maintain confidentiality of sellers and purchasers. Therefore, the selling of these pigs is unregulated and unmonitored, such that in an escape, release, or disease event, there is no paper trail available to trace the animals back to a source once the advertisements are deleted online after the purchases are complete. This study identified six categories of pig types for sale, the most popular by far being pot belly pigs, followed by wild boar. Breeds or types of pigs for sale are often misidentified or mislabelled by sellers (Fig. 1), for this reason genetic testing needs to be employed in order to quantify what is being sold, especially in the case of hybrids. To be required to provide the results of genetic testing on pigs being sold and have available a genetic database for pigs would enable more thorough monitoring of pigs being sold and potentially released, as well as enforcement of proper animal care (i.e., not releasing into the wild or selling under false pretences). Frequently, people with good intentions choose to purchase pigs as pets, under the impression that ‘teacup’ or ‘micro’ pigs will stay small and easy to care for, and now the number of pigs being surrendered are happening much faster than they can be rehomed. (Braich, 2021). Indeed, there are at least 84 farm animal rescues and sanctuaries in Canada, not including Humane Societies or Society’s for the Prevention of Cruelty to Animals (MacDonald, unpublished data).

Most sellers did not disclose if the pigs they were selling were tattooed or tagged. These forms of identification allow trace-back and tracking of individual or groups of pigs, and critical for the swift responses necessary to mitigate disease occurrences in the event of non-native pig release(s) or escape(s) and disease outbreaks (CFIA, 2021). Across Canada all domestic swine are required to be ear tagged and/or tattooed under the PigTrace program and all premises should be recorded. Pig Trace is a national traceability initiative that is mandatory under the federal Health of Animals Regulations (Canadian Pork Council, 2022). Wild pigs are capable of being vectors and/or reservoirs for numerous pathogens and diseases and are a significant risk to the pork industry, wildlife, and public health. As the number and geographic distribution of invasive wild pigs continues to increase rapidly in Canada (Aschim and Brook, 2019; our updated 2022 map), so does the potential risk of exposure and disease transfer to domestic swine.

## 4. Conclusions and Recommendations

Our findings raise important concerns regarding unregulated and undocumented sales of domestic wild boar, pot-bellied pigs, and their hybrids with domestic swine that have important implications for African swine fever preparedness in Canada and the role these sales may play in contributing to the ongoing spread of invasive wild pigs in Canada. While not the focus of this paper, we also observed widespread sales of domestic pigs and this should be examined in more detail. Our literature review has found several studies of backyard small scale swine farming in other countries (Jiménez-Ruiz et al., 2022), but nothing was found for Canada. Further research is needed to better understand the social and ecological implications of small-scale swine farming and the nature of the risks posed by these unregulated farms. Of particular concern is that many of these small pig farms lack expertise on animal husbandry, including adequate fencing and biosecurity. Disease testing of free-ranging domestic swine and these undocumented farms is essential. We recommend that immediate action be taken to regulate online sales of domestic swine in Canada and to ensure that all farms are compliant with the Pig Trace Program. Steps should include further documentation of sales on www.kijiji.ca and other online forums where swine are being sold, with regulations to prevent or at least document these sales. Regulations could include requirements for formalized reporting of any sales, collection of tissue samples for genetic analysis, requirement of ear tags and tattoos, monitoring around domestic farms for escapes and releases of animals, and mandatory disease testing. Canada is one of the largest pork producers in the world, and as such, if ASF were to be introduced into the country, the risk of it quickly spreading and causing catastrophic economic losses is high (Pollock et al., 2021). Education is needed for both producers and those interested in purchasing alternative livestock regarding risks and impacts, including biosecurity and potential contribution to free-ranging invasive wild pigs, as well as risks and impacts of African Swine Fever.

## Acknowledgements

This study was supported by the University of Saskatchewan and the United States Department of Agriculture (USDA). We thank Ruth Aschim for contributing data on the distribution of wild pigs in Canada and Dr. Douglas Clark for thoughtful input on an earlier draft.

## Funding

The background work necessary to complete this study was funded by the United States Department of Agriculture (USDA). The preparation of this paper was funded by the University of Saskatchewan.

## Declaration of Competing Interest

The authors report no declarations of interest.

## Notes

### Competing Interest Statement

The authors have declared no competing interest.

